# Molecular conservation and Differential mutation on ORF3a gene in Indian SARS-CoV2 genomes

**DOI:** 10.1101/2020.05.14.096107

**Authors:** Sk. Sarif Hassan, Pabitra Pal Choudhury, Pallab Basu, Siddhartha Sankar Jana

## Abstract

A global emergency due to the COVID-19 pandemic demands various studies related to genes and genomes of the SARS-CoV2. Among other important proteins, the role of accessory proteins are of immense importance in replication, regulation of infections of the coronavirus in the hosts. The largest accessory proteins in the SARS-CoV2 genome is ORF3a which modulates the host response to the virus infection and consequently it plays an important role in pathogenesis. In this study, an attempt is made to decipher the conservation of nucleotides, dimers, codons and amino acids in the ORF3a genes across thirty two genomes of Indian patients. ORF3a gene possesses single and double point mutations in Indian SARS-CoV2 genomes suggesting the change of SARS-CoV2’s virulence property in Indian patients. We find that the parental origin of the ORF3a gene over the genomes of SARS-CoV2 and Pangolin-CoV is same from the phylogenetic analysis based on conservations of nucleotides and so on. This study highlights the accumulation of mutation on ORF3a in Indian SARS-CoV2 genomes which may provide the designing therapeutic approach against SARS-CoV2.

## 1. Introduction

Since December, 2019, the coronavirus disease (COVID-19) due to the severe acute respiratory syndrome (SARS) originating from Wuhan, China, has been causing a pandemic across the world [1, 2]. The causative virus, SARS-CoV2 is a positive-stranded RNA virus with genome size approximately of 30000 bases. The genome of SARS-CoV2 contains twenty nine open reading frames (ORFs) [3, 4]. Among the twenty nine ORFs, there are sixteen nonstructural proteins (nsps), four structural proteins (E, M, N, S), and six or seven accessory proteins such as ORF3a, ORF6, ORF7a, ORF7b, ORF8 and ORF10 [5, 6, 7]. SARS-CoV2 has been thought to be evolved due to rapid mutation, and recombination with existing other coronavirus in the body. They can alter tissue tropism, cross the species barrier and adopt to different epidemiological situations [8]. Sequence similarity based phylogeny infers that the SARS-CoV2 forms a distinct lineage with Bat-SARS-like coronaviruses that belong to the genus Beta-coronavirus (*β*-CoVs) [9]. The SARS-CoV2 genomes have a significant sequential similarity with percentages 96.3%, 89%, and 82% with bat CoV, SARS-like CoV, and SARS-CoV, respectively, which confirms zoonotic origin of the SARS-CoV2 [10]. There are about 380 amino acid changes from the different proteins of SARS-CoV genomes to the proteins of present SARS-CoV2 genomes as reported so far [11]. The 348, 27 and 5 changes of amino acids occurred in different accessory proteins, S protein and N protein respectively [11]. The accessory proteins have a significant role in virus pathogenesis and these proteins regulate the interferon signalling pathways and the production of pro-inflammatory cytokines [12]. The ORF3a gene which encodes a protein of 274 amino acids, is the second largest sub-genomic RNA in the genome of SARS-CoV [13]. The ORF3a gene encodes a protein with TRAF, ion channel and caveolin binding domain [14]. Mutation in these region alters the NF-kB activation and NLRP3 inflammosome [13]. One of the important features of the ORF3a protein is the presence of a cysteine-rich domain as observed in the SARS-CoV genomes [15]. The ORF3a protein is expressed abundantly in infected and transfected cells, which localizes to intracellular and plasma membranes [16, 17]. It induces apoptosis in transfected and infected cells [18]. In the SARS-CoV genomes, co-mutation between the ORF3a gene and the spike gene exists which suggests that the function of the ORF3a protein correlates with the spike protein [19, 20, 21]. Therefore locating the mutation in ORF3a proteins might lead to understanding the functionality changes in the protein during viral spreading. Till today, no such study has been carried out to look for the existence of ORF3a variation in the Indian patients.

In this present study, we intend to transact the molecular arrangements of nucleotides, dimers, codons and amino acids of the ORF3a gene/protein sequences of SARS-CoV2 of the Indian patients and of CoVs of Bat and Pangolin in order to fetch the evolution connections (if there is any) and similarities and dissimilarities. This study would help to comprehend the effect of non-synonymous mutations in the accessory proteins of the SARS-CoV genomes collected from various geo-locations across the world. In addition, beyond sequence similarity based bioinformatics, this study opens us the hidden conservation of nucleotides, dimers, codons and amino acids over the accessory protein OR3a of three different hosts such as Bat, Pangolin and Human.

### 1.1. Findings on the Dataset

Globally, to this date, among 2385 genomes, we see 118 different mutations in the ORF3a gene. Among these mutations, three changes the size of the gene ORF3a. Out of three changes, one is with deletion of one codon (MT358717-USA: WA), second with deletion of two codons (MT293186-USA: WA) and third with insertion of one codon (MT449656-USA: CA). The rest (115 in total), including accessions from India, contain ORF3a genes of SARS-CoV2 genomes with only point mutations. There are five major genomic groups with sizes (1068, 967, 100, 31, 30), the rest of the groups have sizes in one digit. We name the two largest groups as ORF3a-Type-1 and ORF3a-Type-2. Among them, there is just a difference of one point mutation (G to T) at the 117^*th*^ position of the ORF3a gene across all the 967 SARS-CoV2 genomes. In all the groups, the number of point mutations is found to be at most 4, across the available genome data. The most divergent mutations are often found in the USA. Though 102 different position of ORF3a are globally found, but mutation in three positions which are exclusively in Indian SARS-CoV2 are considered for our study.

As on May 14, 2020, there are thirty two complete genomes viz. MT451874, MT451876, MT451877, MT451878, MT451880, MT451881, MT451882, MT451883, MT451884, MT451885, MT451886, MT451887, MT451888, MT451889, MT451890, MT435079, MT435080, MT435081, MT435082, MT435083, MT435084, MT435085, MT435086, MT415320, MT415321, MT415322, MT415323, MT358637, MT012098 and MT050493 of SARS-CoV2 from Indian patients are available in the NCBI database and that are considered for this present study [22]. Note that, except the genomes MT012098, MT050493 all the other thirty genomes belong to the L-type as per classification made in the article [23]. A set of brief remarks on the accessory protein coding genes across the thirty two genomes from the Indian patients is given in Table 1. The proteins ORF7a, ORF6 and ORF10 are 100% conserved in the thirty-two SARS-CoV2 genomes of Indian origin. However, there are four different types of ORF3a genes that are found based on single-point mutations.

**Table 1:** Accessory proteins coding genes with associated remarks based on the thirty two genomes from India

| Accessory Gene | Remarks Based on the thirty two Indian Genomes |
| --- | --- |
| **ORF3a** | Three single-point mutations ( viz. G to T and C to T) are found in ORF3a gene across the thirty genomes. |
| **ORF6** | 100% identical across all the thirty two genomes. |
| **ORF7b** | 100% identical across all the thirty two genomes. |
| **ORF7a** | 100% identical except in the genome MT435082.<br>From 318th onwards 20 ambiguous base *N* are placed. |
| **ORF10** | 100% identical across all the thirty genomes. |
| **ORF8** | 100% identical except in the genomes MT435081 and MT435082.<br>Note that MT435081 and MT35082 contain the truncated ORF8 gene.<br>In the truncated genes there is a point mutation from *C* to *T*.<br>Note that ORF8 and ORF7a are exactly of same length but it does not have any significant similarity. |

In Indian patients, we found twenty-two ORF3a-Type-1 and seven ORF3a-Type-2 genomes among the thirty-two genomes of the Indian patients. The rest of the two types of mutations (we have seen 2+1=3 genomes) are Indian patients specific and have only one base difference with ORF3a-Type-2 and two bases differences from the 50 ORF3a-Type-1. We named these two Indian groups as ORF3a-Type-3 and ORF3a-Type-4 (refer to Table 2). The nucleotide frequencies, length and some associated remarks of the four types of ORF3a genes of SARS-CoV2 genomes of the Indian patients including the ORF3a genes of the pangolin and Bat CoV are presented in Table 2.

**Table 2:** ORF3a genes across different SARS-CoV2 and CoVs genomes of Pangolin and Bat

| ORF3a/Genome ID | Host | # of A | # of C | # of G | # of T | Length | Remarks |
| --- | --- | --- | --- | --- | --- | --- | --- |
| **ORF3a-Type-1** | Human | 225 | 174 | 153 | 276 | 828 | At 171<sup>th</sup> position, the base is G |
| **ORF3a-Type-2** | Human | 225 | 174 | 152 | 277 | 828 | W.r.t. ORF3a-Type-1 gene, at 171<sup>th</sup> position one mutation G to T occurred. |
| **ORF3a-Type-3** | Human | 225 | 174 | 151 | 278 | 828 | W.r.t. ORF3a-Type-2 gene, at 463<sup>rd</sup> position one mutation G to T occurred. |
| **ORF3a-Type-4** | Human | 225 | 173 | 152 | 278 | 828 | W.r.t. ORF3a-Type-2 gene, at 512<sup>th</sup> position one mutation C to T occurred. |
| **MT040333** | Pangolin | 223 | 175 | 151 | 279 | 828 | The query gene ORF3a |
| **MT040334** | Pangolin | 224 | 173 | 152 | 279 | 828 | 826/828(99%) |
| **MT040335** | Pangolin | 225 | 172 | 152 | 279 | 828 | 825/828(99%) |
| **MT040336** | Pangolin | 224 | 173 | 152 | 279 | 828 | 826/828(99%) |
| **KY417143** | Bat | 223 | 178 | 161 | 263 | 825 | The query gene ORF3a |
| **KY417144** | Bat | 234 | 179 | 152 | 260 | 825 | 749/827(91%) |
| **KY417146** | Bat | 232 | 176 | 156 | 261 | 825 | 751/829(91%) |
| **KY417147** | Bat | 227 | 179 | 158 | 261 | 825 | 807/825(98%) |
| **KY417148** | Bat | 222 | 179 | 162 | 262 | 825 | 821/825(99%) |
| **KY417149** | Bat | 225 | 181 | 158 | 261 | 825 | 795/825(96%) |
| **KY417150** | Bat | 233 | 179 | 153 | 260 | 825 | 748/827(90%) |
| **KY417151** | Bat | 236 | 180 | 151 | 258 | 825 | 745/827(90%) |
| **KY417152** | Bat | 235 | 177 | 150 | 262 | 824 | 748/827(90%) |

In the Table 2, it is found that the length of ORF3a gene of SARS-CoV2 genomes is 828 bases whereas the length of ORF3a gene of SARS-CoV was 825 bases. That is ORF3a gene in SARS-CoV and SARS-CoV2 encode amino acid sequence of length 274 and 275 respectively. Clearly, in the present SARS-CoV2 genomes, the one amino acid E, Glutamic acid is inserted after 240th aa of the ORF3 protein sequence into the ORF3a protein sequence which is shown in the Fig.1.

**Figure 1:**
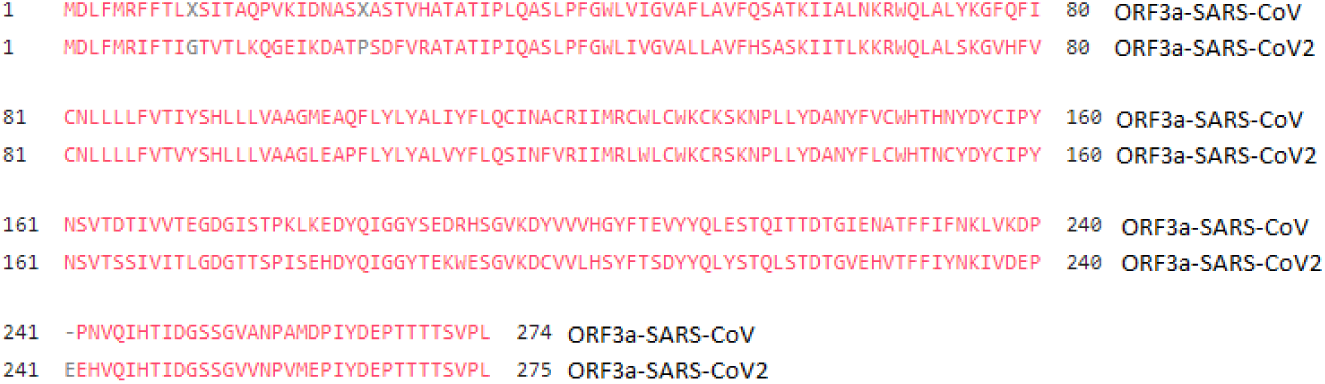
Amino acid Glutamic acid (E) insertion in ORF3a gene of SARS-CoV. Credit: NCBI

The ORF3a protein of the SARS-CoV2 is also blasted (using NCBI-blastp suite) with other ORF3a proteins of Bat and Pangolin CoV. It resulted that the Glutamic acid at the 241^*th*^ position matches with that of Pangolin-CoV which is shown in Fig.2.

**Figure 2:**
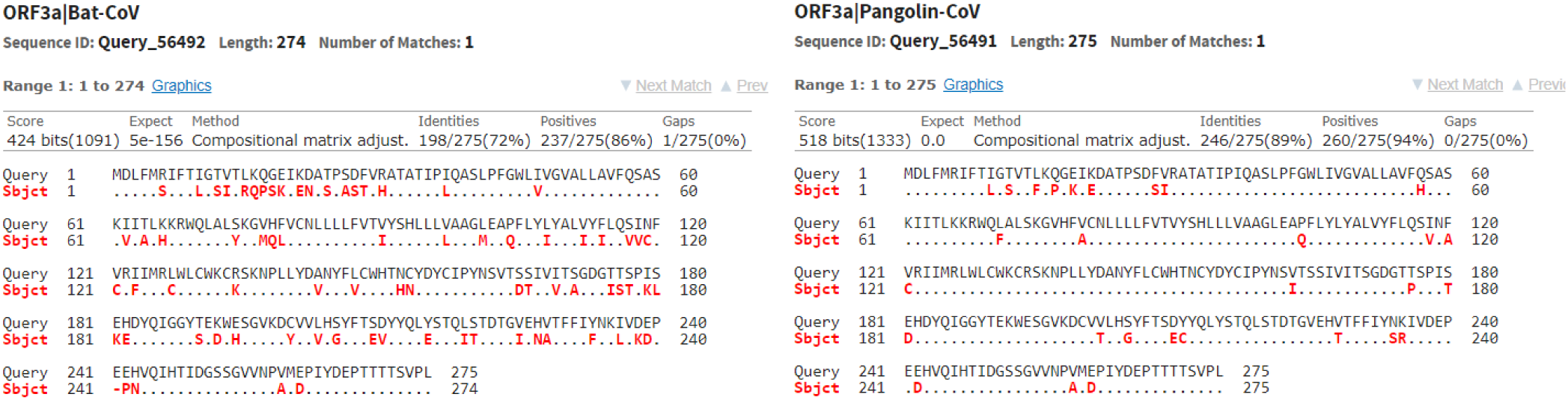
Amino acid sequence alignment of ORF3a across Bat and Pangolin CoV with that of SARS-CoV2. Credit: NCBI

So considering the mutations in ORF3a gene of the SARS-CoV2 genomes of Indian patients, there are four different ORF3a gene sequences of SARS-CoV2 are found and they are referred as ORF3a-Type-1, 2, 3 and 4. These mutations over the gene ORF3a alter the amino acids viz. Q to H, D to Y and S to L), which is schematically presented in the Fig.3.

**Figure 3:**
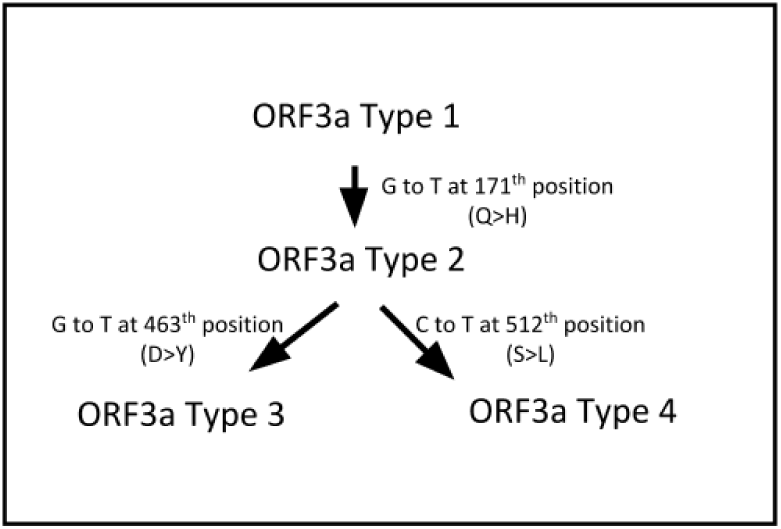
Mutations and associated alternation of amino acids in the four types of ORF3a genes.

The Fig.3 follows that the ORF3a-Type-3 is obtained by two single point mutation (G to T) from the ORF3a-Type-1. Likewise, the ORF3a-Type-4 is achieved by two single point mutations (G to T and C to T) from the ORF3a-Type-1. The genomes which contain the four different types of ORF3a genes of thirty two SARS-CoV2 genomes of the Indian patients are mentioned in Table 3. These data suggest that profiling of mutation on ORF3a genes in Indian patients is different than that of rest of world.

**Table 3:** SARS-CoV2 genomes of 32 Indian patients and their respective type based on the mutation in ORF3a genes

| Accession | Geo_Location | Collection_Date | ORF3a Type | Accession | Geo_Location | Collection_Date | ORF3a Type |
| --- | --- | --- | --- | --- | --- | --- | --- |
| MT457403 | Hyderabad | 2020-03-25 | Type-1 | MT415321 | India | 2020-03-11 | Type-1 |
| MT451874 | Surat | 2020-04-24 | Type-1 | MT415322 | India | 2020-03-16 | Type-1 |
| MT451877 | Surat | 2020-04-26 | Type-1 | MT415323 | India | 2020-03-20 | Type-1 |
| MT451878 | Surat | 2020-04-27 | Type-1 | MT358637 | Rajkot | 2020-04-05 | Type-1 |
| MT451880 | Surat | 2020-04-26 | Type-1 | MT012098 | Kerala State | 2020-01-27 | Type-1 |
| MT451883 | Ahmedabad | 2020-04-26 | Type-1 | MT050493 | Kerala State | 2020-01-31 | Type-1 |
| MT451884 | Ahmedabad | 2020-04-26 | Type-1 | MT457402 | Hyderabad | 2020-03-24 | Type-2 |
| MT451886 | Ahmedabad | 2020-04-26 | Type-1 | MT451876 | India: Surat | 2020-04-26 | Type-2 |
| MT451887 | Ahmedabad | 2020-04-26 | Type-1 | MT451885 | Ahmedabad | 2020-04-26 | Type-2 |
| MT451889 | Ahmedabad | 2020-04-26 | Type-1 | MT451888 | Ahmedabad | 2020-04-26 | Type-2 |
| MT435079 | Ahmedabad | 2020-04-13 | Type-1 | MT435081 | Ahmedabad | 2020-04-13 | Type-2 |
| MT435080 | Ahmedabad | 2020-04-13 | Type-1 | MT435082 | Ahmedabad | 2020-04-13 | Type-2 |
| MT435083 | Ahmedabad | 2020-04-07 | Type-1 | MT435085 | Gandhinagar | 2020-04-22 | Type-2 |
| MT435084 | Ahmedabad | 2020-04-14 | Type-1 | MT451881 | Ahmedabad | 2020-04-26 | Type-3 |
| MT435086 | Mansa | 2020-04-21 | Type-1 | MT451882 | Ahmedabad | 2020-04-26 | Type-3 |
| MT415320 | India | 2020-03-01 | Type-1 | MT451890 | Ahmedabad | 2020-04-26 | Type-4 |

In addition, as the references for establishing any evolutionary connections from the ORF3a gene perspective, ORF3a genes from the four CoV genomes of Pangolin viz. MT040333, MT040334, MT040335 and MT040336 and nine Bat CoV genomes viz. KY417143, KY417144, KY417146, KY417147, KY417148, KY417149, KY417150, KY417151 and KY417152 are considered for the present study. The corresponding phylogeny of the genomes based on sequential similarity of the ORF3a gene is given in the Fig.4.

**Figure 4:**
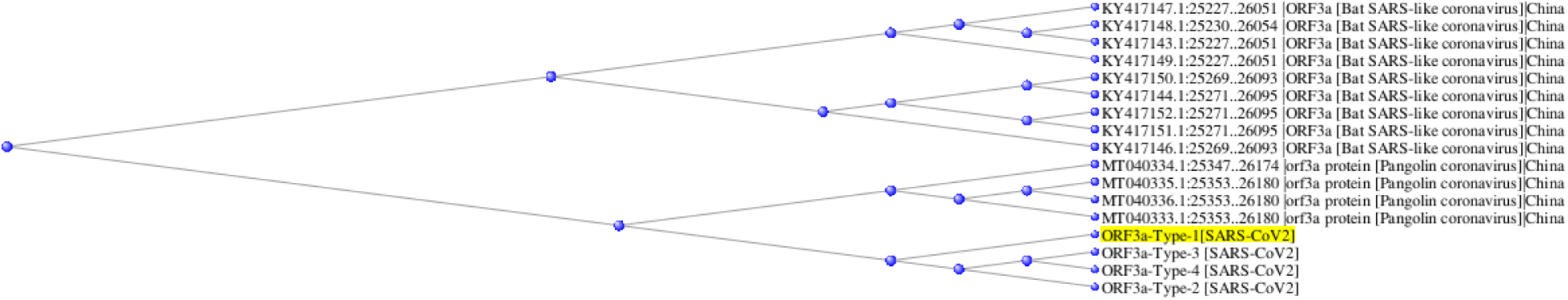
Phylogeny (distance tree) of the thirty genomes based on sequential similarities of the ORF3a genes. Credit: NCBI

The phylogeny shows that the ORF3a genes of CoVs across the three different hosts are mutually placed differently in the distance tree. The phylogeny reports that the ORF3a gene of four types of SARS-CoV2 genomes are sequentially very much closer to that of Pangolin-CoV, than of Bat-CoV. The ORF3a-Type-3 and ORF3a-Type-4 genes are evolved from the ORF3a-Type-2 gene by single point mutations as reported in the phylogeny.

Among 1068 and 967 genomes having mutations of ORF3a-Type-1 and ORF3a-Type-2 respectively, one hundred each such examples of genomes with their respective geo-locations are given in the Table 4 and 5.

**Table 4:** List of accessions and respective geo-locations based on the NCBI blast of the query sequence ORF3a-Type-1 gene.

| Accession | Geo.Location | Accession | Geo.Location | Accession | Geo.Location | Accession | Geo.Location |
| --- | --- | --- | --- | --- | --- | --- | --- |
| MT434758 | India | MT418880 | USA: VA | MT419855 | USA: CA | MT412201 | USA: Michigan |
| MT434759 | India | MT418881 | USA: VA | MT419856 | USA: CA | MT412214 | USA: Michigan |
| MT434760 | India | MT418883 | USA: VA | MT419857 | USA: CA | MT412244 | USA: WA |
| MT434786 | USA: NY | MT418884 | USA: VA | MT419858 | USA: CA | MT412246 | USA: WA |
| MT434796 | USA: NY | MT418893 | USA: VA | MT419859 | USA: CA | MT412248 | USA: WA |
| MT434800 | USA: NY | MT418894 | USA: VA | MT419860 | USA: CA | MT412250 | USA: WA |
| MT434813 | USA: NY | MT419810 | Puerto Rico | MT412134 | China | MT412252 | USA: WA |
| MT435079 | India: Ahmedabad | MT419812 | Puerto Rico | MT412136 | USA: Michigan | MT412253 | USA: WA |
| MT435080 | India: Ahmedabad | MT419815 | Puerto Rico | MT412137 | USA: Michigan | MT412257 | USA: WA |
| MT435083 | India: Ahmedabad | MT419828 | USA: CA | MT412138 | USA: Michigan | MT412261 | USA: WA |
| MT435084 | India: Ahmedabad | MT419829 | USA: CA | MT412139 | USA: Michigan | MT412275 | USA: WA |
| MT435086 | India: Mansa | MT419830 | USA: CA | MT412144 | USA: Michigan | MT412281 | USA |
| MT365028 | Hong Kong | MT419831 | USA: CA | MT412147 | USA: Michigan | MT412290 | USA: WA |
| MT365029 | Hong Kong | MT419832 | USA: CA | MT412157 | USA: Michigan | MT412291 | USA: WA |
| MT365030 | Hong Kong | MT419833 | USA: CA | MT412158 | USA: Michigan | MT412295 | USA: WA |
| MT365031 | Hong Kong | MT419834 | USA: CA | MT412159 | USA: Michigan | MT412302 | USA: CT |
| MT365032 | Hong Kong | MT419835 | USA: CA | MT412167 | USA: Michigan | MT412303 | USA: CT |
| MT428551 | Kazakhstan | MT419837 | USA: CA | MT412172 | USA: Michigan | MT412312 | USA: WA |
| MT428552 | Kazakhstan | MT419839 | USA: CA | MT412173 | USA: Michigan | MT412316 | USA: WA |
| MT428553 | Kazakhstan | MT419841 | USA: CA | MT412174 | USA: Michigan | MT415320 | India |
| MT429187 | USA: Wisconsin | MT419842 | USA: CA | MT412175 | USA: Michigan | MT415321 | India |
| MT429188 | USA: Wisconsin | MT419845 | USA: CA | MT412177 | USA: Michigan | MT415322 | India |
| MT318827 |  | MT419846 | USA: CA | MT412183 | USA: Michigan | MT415323 | India |
| MT270814 | Hong Kong | MT419853 | USA: CA | MT412193 | USA: Michigan | MT415895 | USA: VA |
| MT270815 | Hong Kong | MT419854 | USA: CA | MT412197 | USA: Michigan | MT415896 | USA: VA |

**Table 5:** List of accessions and respective geo-locations based on the NCBI blast of the query sequence ORF3a-Type-2 gene.

| Accession | Geo.Location | Accession | Geo.Location | Accession | Geo.Location | Accession | Geo.Location |
| --- | --- | --- | --- | --- | --- | --- | --- |
| MT434782 | USA: NY | MT434817 | USA: NY | MT419822 | Puerto Rico | MT412216 | USA: Michigan |
| MT434788 | USA: NY | MT435081 | India: Ahmedabad | MT419851 | USA: CA | MT412217 | USA: Michigan |
| MT434789 | USA: NY | MT435082 | India: Ahmedabad | MT412187 | USA: Michigan | MT412218 | USA: Michigan |
| MT434790 | USA: NY | MT435085 | India: Gandhinagar | MT412188 | USA: Michigan | MT412219 | USA: Michigan |
| MT434791 | USA: NY | MT429183 | USA: Wisconsin | MT412189 | USA: Michigan | MT412220 | USA: Michigan |
| MT434792 | USA: NY | MT429184 | USA: Wisconsin | MT412190 | USA: Michigan | MT412221 | USA: Michigan |
| MT434793 | USA: NY | MT429185 | USA: Wisconsin | MT412191 | USA: Michigan | MT412222 | USA: Michigan |
| MT434794 | USA: NY | MT429186 | USA: Wisconsin | MT412192 | USA: Michigan | MT412223 | USA: Michigan |
| MT434795 | USA: NY | MT429189 | USA: Wisconsin | MT412194 | USA: Michigan | MT412224 | USA: Michigan |
| MT434797 | USA: NY | MT429190 | USA: Wisconsin | MT412195 | USA: Michigan | MT415894 | USA: VA |
| MT434798 | USA: NY | MT429191 | USA: Wisconsin | MT412196 | USA: Michigan | MT415897 | USA: VA |
| MT434799 | USA: NY | MT432195 | USA: Louisiana | MT412198 | USA: Michigan | MT415898 | USA: VA |
| MT434801 | USA: NY | MT422806 | USA: FL | MT412199 | USA: Michigan | MT415899 | USA: VA |
| MT434802 | USA: NY | MT422807 | USA: FL | MT412200 | USA: Michigan | MT415900 | USA: VA |
| MT434803 | USA: NY | MT418889 | USA: VA | MT412202 | USA: Michigan | MT415901 | USA: VA |
| MT434804 | USA: NY | MT418890 | USA: VA | MT412203 | USA: Michigan | MT415902 | USA: VA |
| MT434805 | USA: NY | MT418891 | USA: VA | MT412204 | USA: Michigan | MT415903 | USA: VA |
| MT434806 | USA: NY | MT418892 | USA: VA | MT412205 | USA: Michigan | MT415904 | USA: VA |
| MT434808 | USA: NY | MT419811 | Puerto Rico | MT412206 | USA: Michigan | MT415905 | USA: VA |
| MT434809 | USA: NY | MT419814 | Puerto Rico | MT412207 | USA: Michigan | MT415906 | USA: VA |
| MT434810 | USA: NY | MT419817 | Puerto Rico | MT412209 | USA: Michigan | MT415907 | USA: VA |
| MT434811 | USA: NY | MT419818 | Puerto Rico | MT412211 | USA: Michigan | MT415908 | USA: VA |
| MT434812 | USA: NY | MT419819 | Puerto Rico | MT412212 | USA: Michigan | MT415909 | USA: VA |
| MT434815 | USA: NY | MT419820 | Puerto Rico | MT412213 | USA: Michigan | MT415910 | USA: VA |
| MT434816 | USA: NY | MT419821 | Puerto Rico | MT412215 | USA: Illinois | MT415912 | USA: VA |

So these two types of ORF3a gene having one base difference belong to a large class of SARS-CoV2 genomes across different geo-locations as shown in Table 4 and 5. It is noted that the NCBI blast results no genome from China having 100% similarity with the ORF3a-Type-2 gene. That is the one point mutation (G to T) in the ORF3a-Type-2 gene that has happened outside the patients of China. It is worth mentioning that the OF3a-Type-3 and ORF3a-Type-4 genes were blasted in the NCBI database and do not find any 100% similar sequence with 100% query coverage. Hence these two type of mutations in the gene ORF3a are unique in Indian patients.

### 1.2. Methods

In order to determine the molecular level conservations and descriptions of the ORF3a genes across different hosts as mentioned, some methods are discussed [24, 25, 26, 27, 28, 29], which would be subsequently used.

#### Nucleotide Conservation Shannon Entropy

Shannon entropy is a measure of the amount of information (measure of uncertainty). Conservation of each of the four nucleotides has been determined using Shannon entropy [30, 31]. Note that it is assumed *log*_*b*_(0) = 0 for smooth calculation of the SE. For a given sequence of length *l*, the conservation SE (Conv SE) is calculated as follows:

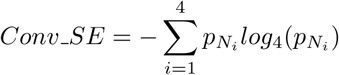

where 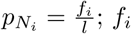 represents the occurrence frequency of a nucleotide *N*_*i*_ in the given sequence.

#### Dimer Conservation Shannon Entropy

The conservation of usages of all possible sixteen dimers (words of length two consisting letters from the set {*A, T, C, G*}) has been determined using Shannon entropy as follows. For a given sequence of length *l*, the conservation of dimers (Dim SE) is calculated as follows:

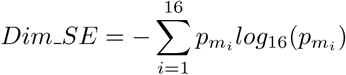

where 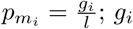 represents the number of occurrences of a dimer *m*_*i*_ in the given sequence.

#### Codon Conservation Shannon Entropy

The conservation of usages of all possible sixty four codons has been determined using Shannon entropy as follows [32]. For a given sequence of length *l*, the conservation of codons (Codon SE) is calculated as follows:

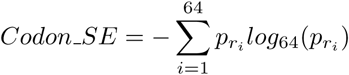

where 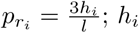 represents the number of occurrences of a codon *r*_*i*_ in the given sequence.

#### Amino Acid Conservation Shannon Entropy

The conservation of twenty amino acids usages across the primary protein sequence encoded by the gene ORF3a has been determined using Shannon entropy as follows. For a given amino acid sequence corresponding to a RNA sequence (ORF3a gene) of length *l*, the conservation of codons (AA SE) is calculated as follows:

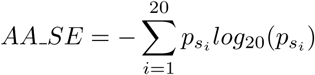

where 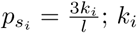 represents the number of occurrences of an amino acid *s*_*i*_ in the given sequence.

In addition to the different conservation SEs, some basic derivative features such as nucleotide frequency and density, frequency of all possible sixteen dimers, frequency of codon usages, frequency of amino acids in the protein sequence encoded by the ORF3a gene, GC content, pyrimidine density are obtained for a given ORF3a gene sequence [24, 26]. It is worth noting that the first positive frame has been considered to determine codons and double nucleotides over a given gene.

## 2. Results

For each of the seventeen different ORF3a genes (including the genomes of SARS-CoV2, Pangolin and Bat CoV) a feature vector is defined which comprises the nucleotides, dimers, codons and amino acids frequencies and associated conservations in the ORF3a genes. Based on these feature vectors corresponding to each of the seventeen sequences, a nearest neighbourhood joining phylogeny is built up for each of the molecular conservations of nucleotides, dimers, codon and amino acids.

### 2.1. Frequency and Conservation of nucleotides over ORF3a Gene

The counts of the nucleotide bases, length, GC content and pyrimidine density (py density) and the conservation Shannon entropy (ConV SE) of the seventeen ORF3a genes across three different hosts are tabulated in Table 6.

**Table 6:** Molecular descriptions of the gene ORF3a across different hosts

| ORF3a/Genome ID | Den A | Den C | Den G | Den T | GC Content | Py Density | Conv_SE |
| --- | --- | --- | --- | --- | --- | --- | --- |
| **ORF3a-Type-1** | 0.2717 | 0.2101 | 0.1848 | 0.3333 | 39.4928 | 54.3478 | 0.9811 |
| **ORF3a-Type-2** | 0.2717 | 0.2101 | 0.1836 | 0.3345 | 39.3720 | 54.4686 | 0.9806 |
| **ORF3a-Type-3** | 0.2717 | 0.2101 | 0.1824 | 0.3357 | 39.2512 | 54.5894 | 0.9801 |
| **ORF3a-Type-4** | 0.2717 | 0.2089 | 0.1836 | 0.3357 | 39.2512 | 54.4686 | 0.9802 |
| **MT040333** | 0.2693 | 0.2114 | 0.1824 | 0.3370 | 39.3720 | 54.8309 | 0.9801 |
| **MT040334** | 0.2705 | 0.2089 | 0.1836 | 0.3370 | 39.2512 | 54.5894 | 0.9800 |
| **MT040335** | 0.2717 | 0.2077 | 0.1836 | 0.3370 | 39.1304 | 54.4686 | 0.9798 |
| **MT040336** | 0.2705 | 0.2089 | 0.1836 | 0.3370 | 39.2512 | 54.5894 | 0.9800 |
| **KY417143** | 0.2703 | 0.2158 | 0.1952 | 0.3188 | 41.0909 | 53.4545 | 0.9867 |
| **KY417144** | 0.2836 | 0.2170 | 0.1842 | 0.3152 | 40.1212 | 53.2121 | 0.9843 |
| **KY417146** | 0.2812 | 0.2133 | 0.1891 | 0.3164 | 40.2424 | 52.9697 | 0.9849 |
| **KY417147** | 0.2752 | 0.2170 | 0.1915 | 0.3164 | 40.8485 | 53.3333 | 0.9862 |
| **KY417148** | 0.2691 | 0.2170 | 0.1964 | 0.3176 | 41.3333 | 53.4545 | 0.9873 |
| **KY417149** | 0.2727 | 0.2194 | 0.1915 | 0.3164 | 41.0909 | 53.5758 | 0.9866 |
| **KY417150** | 0.2824 | 0.2170 | 0.1855 | 0.3152 | 40.2424 | 53.2121 | 0.9846 |
| **KY417151** | 0.2861 | 0.2182 | 0.1830 | 0.3127 | 40.1212 | 53.0909 | 0.9843 |
| **KY417152** | 0.2852 | 0.2148 | 0.1820 | 0.3180 | 39.6845 | 53.2767 | 0.9829 |

The density of each nucleotide bases across the seventeen ORF3a genes are plotted in the Fig.5.

**Figure 5:**
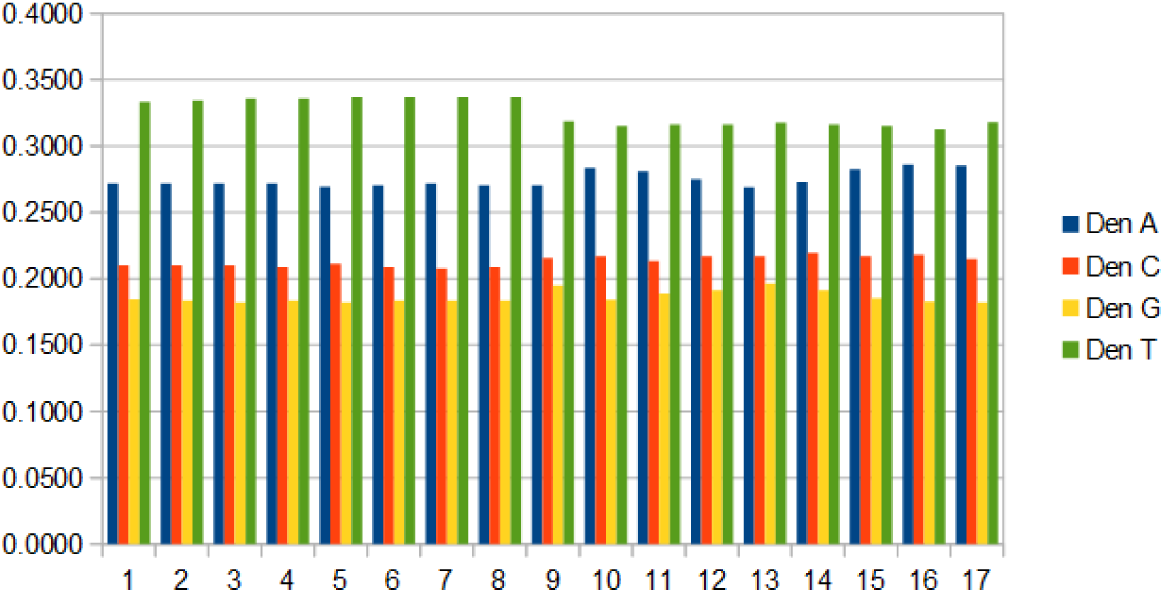
Nucleotide density of four bases across the seventeen ORF3a genes. The numbers 1, 2, 3, … denote the ORF3a gene/Genome ID from the top to bottom of the first column of Table 6, respectively.

In each ORF3a gene the density of T is maximum and G is minimum. Also it is noted the density of C dominates that of G over all the ORF3a genes of three different hosts. The ORF3a gene are pyrimidine-rich with percentage approximately 53% across different genomes as mentioned in the Table 6. Also the ORF3a possesses highest GC content across the Bat CoV genomes and which is ranging from 39.68% to 41.34%. After a single mutation, the GC content of ORF3a-Type-2 is slightly reduced to 39.5% from that of the ORF3a-Type-1 gene. The GC content of Pangolin CoVs is turned out to be minimum and that is 39.13%. The GC content of ORF3a-Type-2 gene and ORF3a of MT040333 is identical though the density of G and C are slightly different in the respective sequences. The ORF3a genes across fifteen different genomes of CoV of the three hosts are highly conserved with equally likely presence of the four nucleotide bases as the Conv SE for all the genes are turned out to be approximately 1.

Based on features of the ORF3a gene across the seventeen CoV genomes, as mentioned in the Table 6, a phylogeny has been developed as shown in Fig.6.

**Figure 6:**
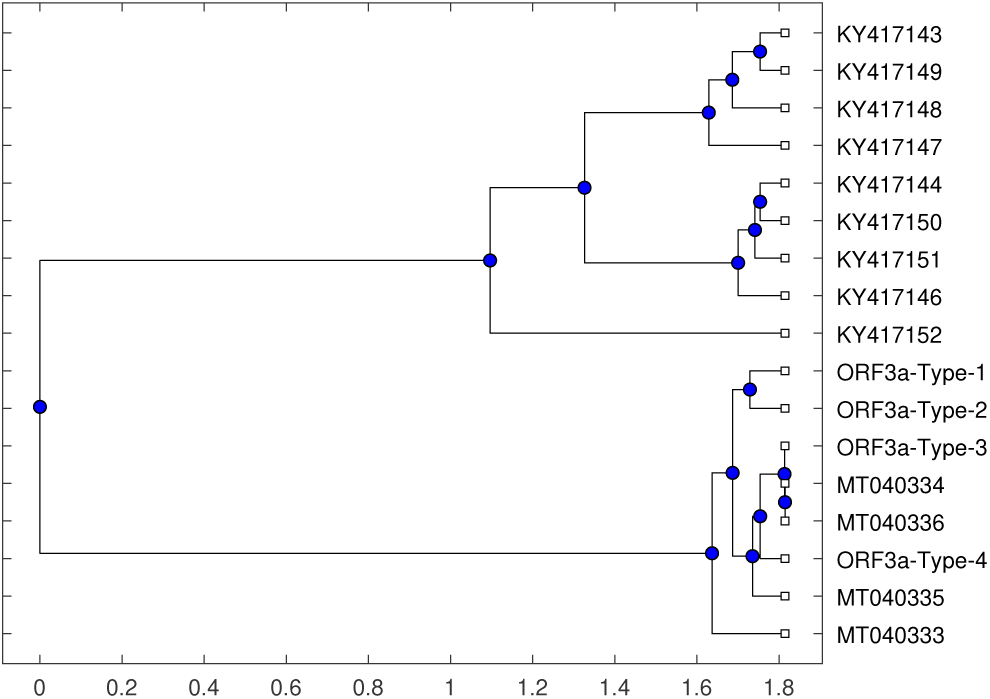
Phylogenetic relationships among the seventeen CoV genomes based on the densities of nucleotides of ORF3a gene

The phylogeny depicts that the ORF3a-Type-1 and ORF3a-Type-2 gene of the SARS-CoV2 genomes of the Indian patients are very close to each other (belong to the 4th level of the tree). At the 6th level of the phylogenetic tree, the ORF3a-Type-3 and that of the genomes MT040334 and MT040336 of CoV of Pangolin belong and naturally they are co-evolved from the previous evolutionary levels of the tree. The ORF3a-Type-4 gene and ORF3a of the Bat CoV genome MT040335 belong to the binary branch of 4th level of the phylogenetic tree. It is also inferred from the Fig.3 that the ORF3a genes of four types of SARS-CoV2 and CoV-Pangolin are evolved from the ORF3a gene of the Pangolin CoV genome MT040333. On the other side, ORF3a gene of Bat CoV are distantly placed in the tree. Among the nine genomes of Bat CoV, the pair of genomes {*KY* 417143, *KY* 417149} and {*KY* 417144, *KY* 417150} are the nearest as they belong to the sixth level of the tree.

### 2.2. Frequency and Conservation of dimers over ORF3a Gene

All possible words consisting letters from the set {*A, T, C, G*} of length two are commonly known as dimers. The frequency of dimers and conservation Shannon entropy of dimers (Dim SE) over the seventeen ORF3a genes across various genomes of CoV are presented in the Table 7. Also a graphical representation of the frequency of the dimers of four types (dimers start with A, T, C and G) are given in Fig.7.

**Figure 7:**
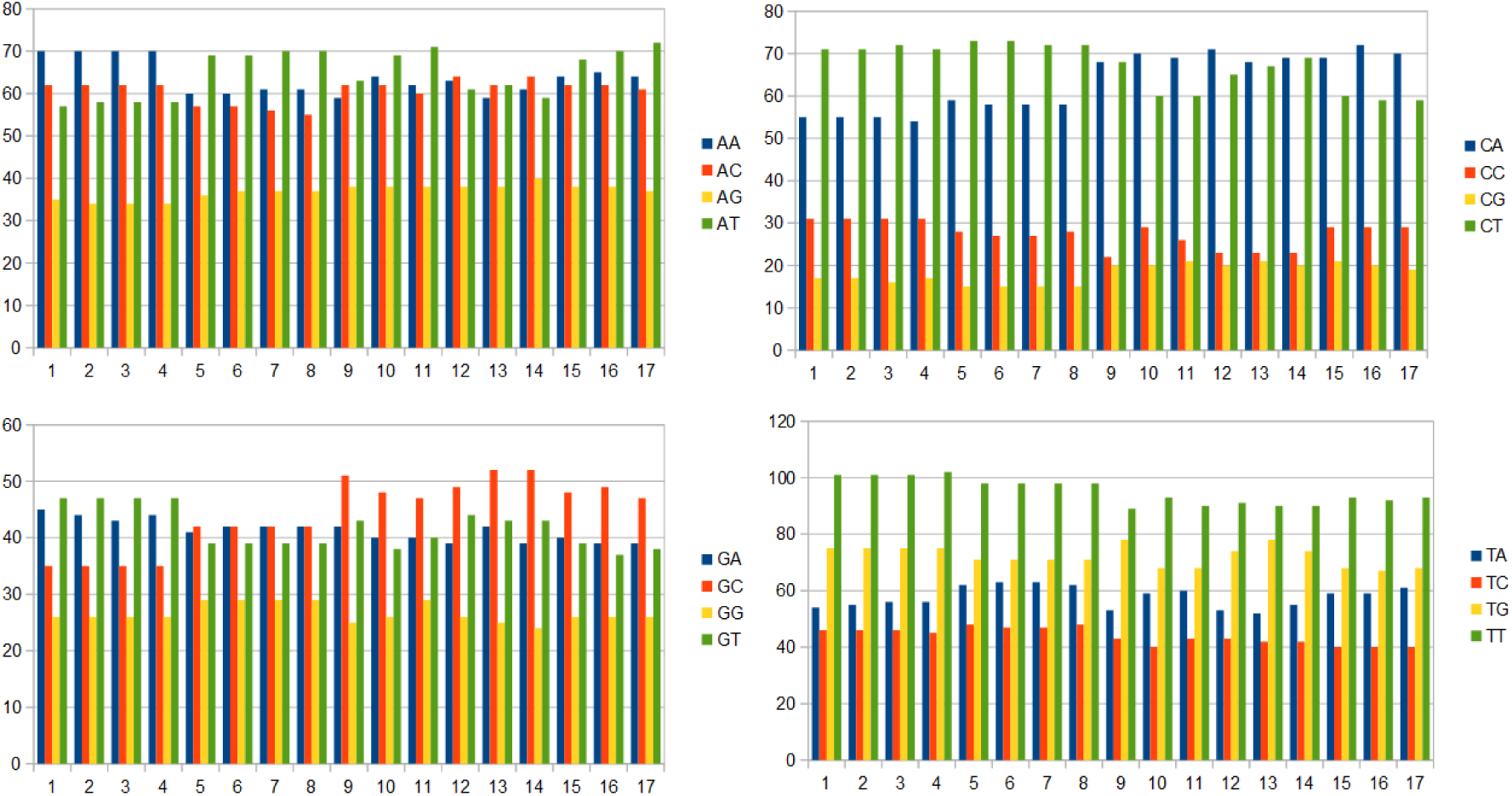
Bar-plot of the frequencies of dimers of ORF3a genes

**Table 7:** Frequency of dimers of the gene ORF3a and associated dimer conservation Shannon entropy

| ORF3a/Genome ID | AA | AC | AG | AT | CA | CC | CG | CT | GA | GC | GG | GT | TA | TC | TG | TT | Dim_SE |
| --- | --- | --- | --- | --- | --- | --- | --- | --- | --- | --- | --- | --- | --- | --- | --- | --- | --- |
| ORF3a-Type-1 | 70 | 62 | 35 | 57 | 55 | 31 | 17 | 71 | 45 | 35 | 26 | 47 | 54 | 46 | 75 | 101 | 0.9705 |
| ORF3a-Type-2 | 70 | 62 | 34 | 58 | 55 | 31 | 17 | 71 | 44 | 35 | 26 | 47 | 55 | 46 | 75 | 101 | 0.9702 |
| ORF3a-Type-3 | 70 | 62 | 34 | 58 | 55 | 31 | 16 | 72 | 43 | 35 | 26 | 47 | 56 | 46 | 75 | 101 | 0.9694 |
| ORF3a-Type-4 | 70 | 62 | 34 | 58 | 54 | 31 | 17 | 71 | 44 | 35 | 26 | 47 | 56 | 45 | 75 | 102 | 0.9698 |
| MT040333 | 60 | 57 | 36 | 69 | 59 | 28 | 15 | 73 | 41 | 42 | 29 | 39 | 62 | 48 | 71 | 98 | 0.9706 |
| MT040334 | 60 | 57 | 37 | 69 | 58 | 27 | 15 | 73 | 42 | 42 | 29 | 39 | 63 | 47 | 71 | 98 | 0.981 |
| MT040335 | 61 | 56 | 37 | 70 | 58 | 27 | 15 | 72 | 42 | 42 | 29 | 39 | 63 | 47 | 71 | 98 | 0.9705 |
| MT040336 | 61 | 55 | 37 | 70 | 58 | 28 | 15 | 72 | 42 | 42 | 29 | 39 | 62 | 48 | 71 | 98 | 0.9709 |
| KY417143 | 59 | 62 | 38 | 63 | 68 | 22 | 20 | 68 | 42 | 51 | 25 | 43 | 53 | 43 | 78 | 89 | 0.9726 |
| KY417144 | 64 | 62 | 38 | 69 | 70 | 29 | 20 | 60 | 40 | 48 | 26 | 38 | 59 | 40 | 68 | 93 | 0.9738 |
| KY417146 | 62 | 60 | 38 | 71 | 69 | 26 | 21 | 60 | 40 | 47 | 29 | 40 | 60 | 43 | 68 | 90 | 0.9756 |
| KY417147 | 63 | 64 | 38 | 61 | 71 | 23 | 20 | 65 | 39 | 49 | 26 | 44 | 53 | 43 | 74 | 91 | 0.9729 |
| KY417148 | 59 | 62 | 38 | 62 | 68 | 23 | 21 | 67 | 42 | 52 | 25 | 43 | 52 | 42 | 78 | 90 | 0.9733 |
| KY417149 | 61 | 64 | 40 | 59 | 69 | 23 | 20 | 69 | 39 | 52 | 24 | 43 | 55 | 42 | 74 | 90 | 0.9727 |
| KY417150 | 64 | 62 | 38 | 68 | 69 | 29 | 21 | 60 | 40 | 48 | 26 | 39 | 59 | 40 | 68 | 93 | 0.9746 |
| KY417151 | 65 | 62 | 38 | 70 | 72 | 29 | 20 | 59 | 39 | 49 | 26 | 37 | 59 | 40 | 67 | 92 | 0.9735 |
| KY417152 | 64 | 61 | 37 | 72 | 70 | 29 | 19 | 59 | 39 | 47 | 26 | 38 | 61 | 40 | 68 | 93 | 0.9727 |

From the Fig.7, it is noticed that frequency of the dimers starting with the letter T is the highest over the gene ORF3a across the seventeen distinct genomes. The frequency decreases over the dimers with the first letter A, C and G respectively. The dimers TT and CG attain maximum and minimum frequency over the ORF3a gene across the fifteen genomes. In all the four types of ORF3a genes the frequencies of the dimers AG, AT, CA, CG, CT, GA, TA, TC and TT are varying as observed in the Table 7. The frequency of usages of most of the dimers in the ORF3a genes of four types dominates that of the Pangolin and Bat CoVs. The Dim SE follows that the ORF3a genes across all the genomes are conserved with all sixteen dimers with nearly equal probability of occurrences. The Dim SE of the ORF3a-Type-1 and ORF3a of the genome MT040335 of Pangolin-CoV are identical although the frequency of respective dimers are different. It is noted that all the dimers are equally likely to appear and conserved in the ORF3a-Type-3 and ORF3a-Type-4 genes.

Based on the frequency of dimers across the ORF3a genes over the genomes the following phylogeny is made in Fig.8.

**Figure 8:**
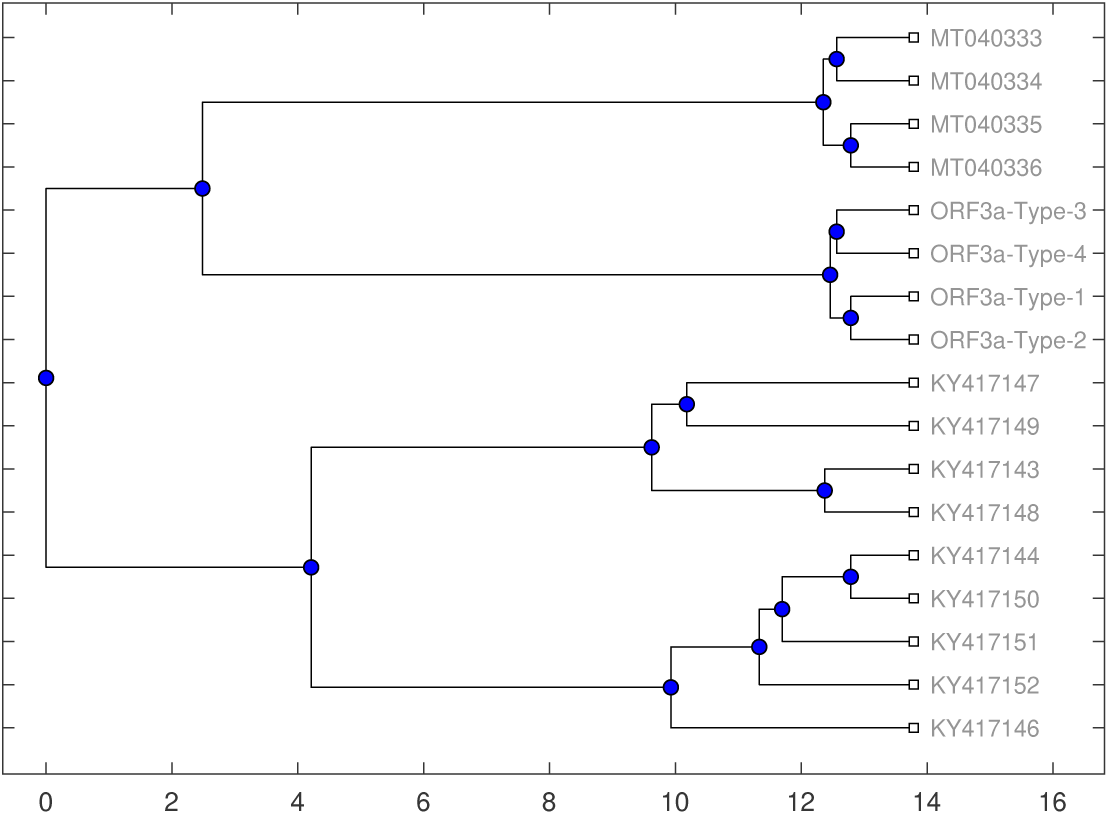
Phylogenetic relationships among the seventeen CoV genomes based on the frequency of dimers of ORF3a genes

The phylogeny based on the frequency distribution of the dimers over the ORF3a genes across various genomes of different hosts follows that ORF3a genes of SARS-CoV2 genomes of Indian patients and genomes of Pangolin-COV are co-evolved by belonging into the same level of the tree. In the other branch of the phylogenetic tree ORF3a genes of the Bat CoV are placed and among them the genomes KY417144 and KY417150 are the nearest based on the dimers usages over the gene ORF3a as found in the Fig.8.

### 2.3. Codon conservations and associated Descriptions of ORF3a Gene

The frequency usages of all the codons over the ORF3a genes across the SARS-CoV2 genomes of Indian patients including genomes of Pangolin and Bat CoVs are given in Table 8. All the twenty amino acids are present in the protein sequence of ORF3a although It is observed that the codons CCC, CGA, GGG, TAG and TGA are thoroughly absent from the ORF3a genes across all the genomes. The ORF3a genes of SARS-CoV2 genomes of the Indian patients as well as of Pangolin CoV contain one CGC while that of the Bat CoV do not contain the codon CGC. This presence of the codon CGC (codes Arginine) deviates the ORF3a gene of SARS-CoV2 and Pangolin CoV from that of the Bat-CoV. In contrast, ORF3a genes of the genomes of Bat-CoV contain the codon GCG (encode Alanine) while the ORF3a genes of four types of SARS-CoV2 genomes do not contain it. It is found that the frequency of GAG, GTG in ORF3a genes of Bat-CoV dominates that of the other two host genomes. The most preferred stop codon across all the ORF3a genes of various CoV genomes is TAA. The most frequently used codon ATT and GTT which encode Isoleucine and Valine respectively in ORF3a across all the observed genomes. The ORF3a genes possess clearly codon biases in encoding the various amino acids as evident from the codon frequency usages.

**Table 8:**
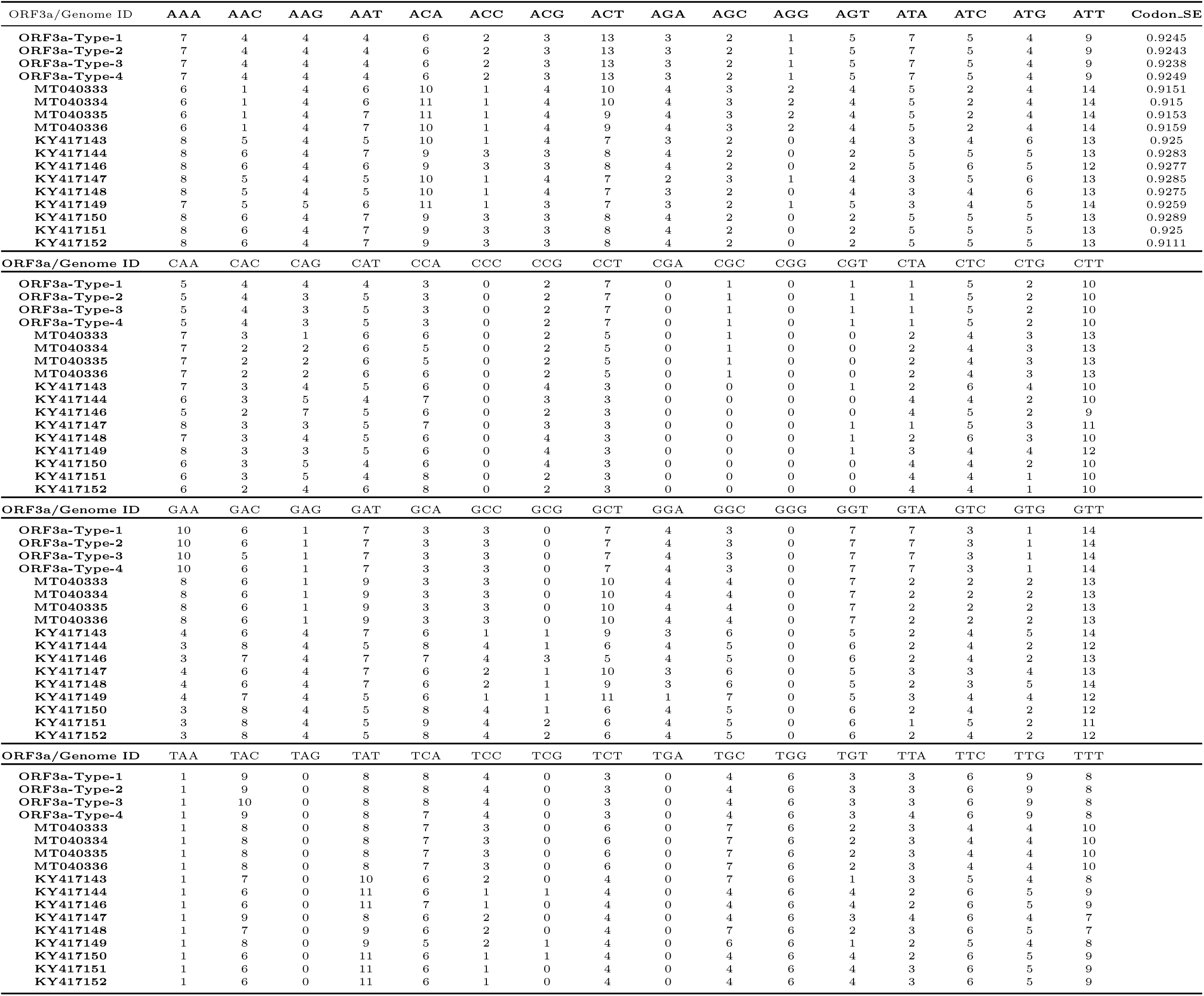
Frequency of codon usages over the gene ORF3a across the seventeen CoV genomes

Over the seventeen different genomes of SARS-CoV2, Pangolin and Bat, the codons are not as conserved as the nucleotides and dimers were in the ORF3a gene due to the codon biases. The Codon SE of ORF3a genes across the genomes are ranging from 0.9111 to 0.9289 and this emerges to a certain degree of uncertainty of codon conservation over the gene.

The following phylogeny of the seven genomes is made by using the frequency of codon usages over the gene ORF3a, as shown in Fig.9.

**Figure 9:**
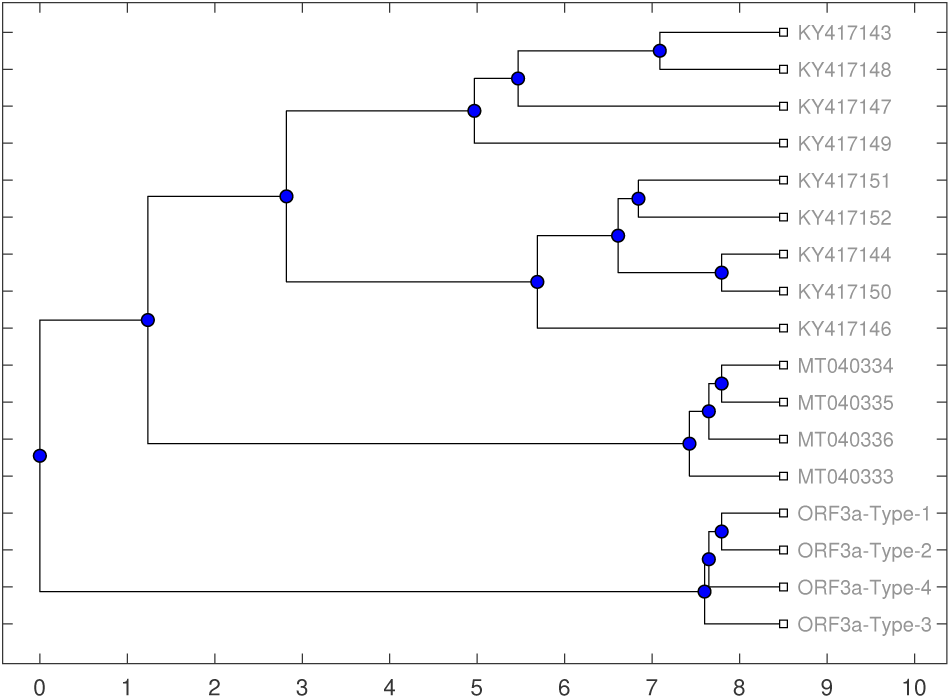
Phylogenetic relationships among the seventeen CoV genomes based on the frequency of codon usages in ORF3a gene across fifteen genomes

**Figure 10:**
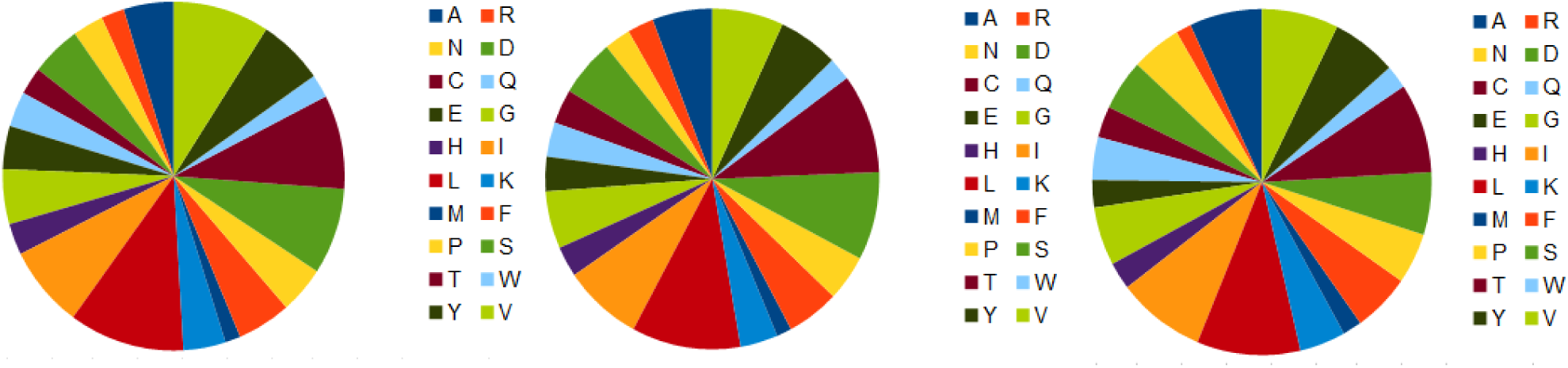
Frequency distribution of amino acids over the ORF3a genes of SARS-CoV2 genomes of the Indian patients, Pangolin-CoV and Bat CoV from left to right.

Based on frequency of codon usages and conservation of codon in the ORF3a genes, the four types of SARS-CoV2 genomes of the Indian patients are distantly placed from the Pangolin and Bat CoVs as chalked out in the phylogenetic tree. The closest distribution of codons in the gene ORF3a over the pair of genomes KY417143 and KY417148 of Bat-CoV is noted. This phylogeny in the Fig.9 depicts that the ORF3a gene of genomes of the Indian patients and that of Bat CoV are co-evolved from the same origin.

### 2.4. Amino acids conservations and associated Descriptions of ORF3a Gene

The frequency of amino acids over the gene ORF3a across the genome of Indian patients, Pangolin and Bat are presented in the Table 9. All the twenty amino acids are present over the gene ORF3a across all the genomes and it is turned out that the ORF3a protein is Luicine-rich with percentage approximately 10%. It is worth mentioning that the ORF3a gene of SARs-CoV genomes were cystine rich []. The frequency of the amino acids Methionine and Arginine are the lowest among all over the ORF3a genes across the genomes.

**Table 9:** Amino acids frequencies over the ORF3a protein sequence across the seventeen genomes

| ORF3a/Genome ID | A | R | N | D | C | Q | E | G | H | I | L | K | M | F | P | S | T | W | Y | V | AA_SE |
| --- | --- | --- | --- | --- | --- | --- | --- | --- | --- | --- | --- | --- | --- | --- | --- | --- | --- | --- | --- | --- | --- |
| ORF3a-Type-1 | 13 | 6 | 8 | 13 | 7 | 9 | 11 | 14 | 8 | 21 | 30 | 11 | 4 | 14 | 12 | 22 | 24 | 6 | 17 | 25 | 0.9553 |
| ORF3a-Type-2 | 13 | 6 | 8 | 13 | 7 | 8 | 11 | 14 | 9 | 21 | 30 | 11 | 4 | 14 | 12 | 22 | 24 | 6 | 17 | 25 | 0.9553 |
| ORF3a-Type-3 | 13 | 6 | 8 | 12 | 7 | 8 | 11 | 14 | 9 | 21 | 30 | 11 | 4 | 14 | 12 | 22 | 24 | 6 | 18 | 25 | 0.9549 |
| ORF3a-Type-4 | 13 | 6 | 8 | 13 | 7 | 8 | 11 | 14 | 9 | 21 | 31 | 11 | 4 | 14 | 12 | 21 | 24 | 6 | 17 | 25 | 0.9549 |
| MT040333 | 16 | 7 | 7 | 15 | 9 | 8 | 9 | 15 | 9 | 21 | 29 | 10 | 4 | 14 | 13 | 23 | 25 | 6 | 16 | 19 | 0.9587 |
| MT040334 | 16 | 7 | 7 | 15 | 9 | 9 | 9 | 15 | 8 | 21 | 29 | 10 | 4 | 14 | 12 | 23 | 26 | 6 | 16 | 19 | 0.9579 |
| MT040335 | 16 | 7 | 8 | 15 | 9 | 9 | 9 | 15 | 8 | 21 | 29 | 10 | 4 | 14 | 12 | 23 | 25 | 6 | 16 | 19 | 0.9594 |
| MT040336 | 16 | 7 | 8 | 15 | 9 | 9 | 9 | 15 | 8 | 21 | 29 | 10 | 4 | 14 | 13 | 23 | 24 | 6 | 16 | 19 | 0.9602 |
| KY417143 | 17 | 4 | 10 | 13 | 8 | 11 | 8 | 14 | 8 | 20 | 29 | 12 | 6 | 13 | 13 | 18 | 22 | 6 | 17 | 25 | 0.9607 |
| KY417144 | 19 | 4 | 13 | 13 | 8 | 11 | 7 | 15 | 7 | 23 | 27 | 12 | 5 | 15 | 13 | 16 | 23 | 6 | 17 | 20 | 0.961 |
| KY417146 | 19 | 4 | 12 | 14 | 8 | 12 | 7 | 15 | 7 | 23 | 27 | 12 | 5 | 15 | 11 | 16 | 23 | 6 | 17 | 21 | 0.9603 |
| KY417147 | 19 | 4 | 10 | 13 | 7 | 11 | 8 | 14 | 8 | 21 | 28 | 12 | 6 | 13 | 13 | 19 | 22 | 6 | 17 | 23 | 0.9607 |
| KY417148 | 18 | 4 | 10 | 13 | 9 | 11 | 8 | 14 | 8 | 20 | 29 | 12 | 6 | 13 | 13 | 18 | 22 | 6 | 16 | 24 | 0.9619 |
| KY417149 | 19 | 5 | 11 | 12 | 7 | 11 | 8 | 13 | 8 | 21 | 29 | 12 | 5 | 13 | 13 | 19 | 22 | 6 | 17 | 23 | 0.9602 |
| KY417150 | 19 | 4 | 13 | 13 | 8 | 11 | 7 | 15 | 7 | 23 | 27 | 12 | 5 | 15 | 13 | 16 | 23 | 6 | 17 | 20 | 0.961 |
| KY417151 | 21 | 4 | 13 | 13 | 8 | 11 | 7 | 15 | 7 | 23 | 27 | 12 | 5 | 15 | 13 | 15 | 23 | 6 | 17 | 19 | 0.9607 |
| KY417152 | 20 | 4 | 13 | 13 | 8 | 10 | 7 | 15 | 8 | 23 | 27 | 12 | 5 | 15 | 13 | 15 | 23 | 6 | 17 | 20 | 0.9619 |

In the ORF3a gene of Type-1 and Type-2 the frequency of Glutamine and Histidine are altered from 9 to 8 and 8 to 9 respectively. The frequencies of Aspertic acid (D), Leucine (L) are 12 and 30 respectively in the ORF3a-Type-3 gene while those of D and L are 13 and 31 in the ORF3a-Type-4 gene of SARS-CoV2 genomes of the Indian patients. The frequencies of Serine and Tyrosine are increased by 1 in ORF3a while it switches from the Type-3 to Type-4 of SARS-CoV2 genomes of Indian patients.

A typical frequency distribution of amino acids in ORF3a genes across the seventeen genomes are presented in Fig.7. The frequencies of amino acids Isoleucine, Methionine, Phenylalanine and Tryptophan are invariant in ORF3a gene across the SARS-CoV2 and Pangolin-CoV genomes among three hosts.

The AA SE follows that the conservation of amino acids of ORF3a over the genome of Indian patients is invariant under mutation. It is noted that the ORF3a genes over the CoV genomes of Pangolin and Bat possess higher conservation of amino acids than that of SARS-CoV2 genomes of the Indian patients. ORF3a gene over the genomes KY417148 and KY417152 attain the highest amount of amino acid conservations as found in the Table 9.

Based on the frequency distribution of amino acids the following phylogeny (Fig.11) of the seventeen genomes are established. At the fifth level of the phylogenetic tree the pairs of genomes {*ORF* 3*a* − *Type* − 1, *ORF* 3*a* − *Type* − 2}, {*MT* 040335, *MT* 040336} and {*KY* 417144, *KY* 417150} belong as leaf nodes and this imply the co-evolution of the ORF3a gene from the same parental origin.

**Figure 11:**
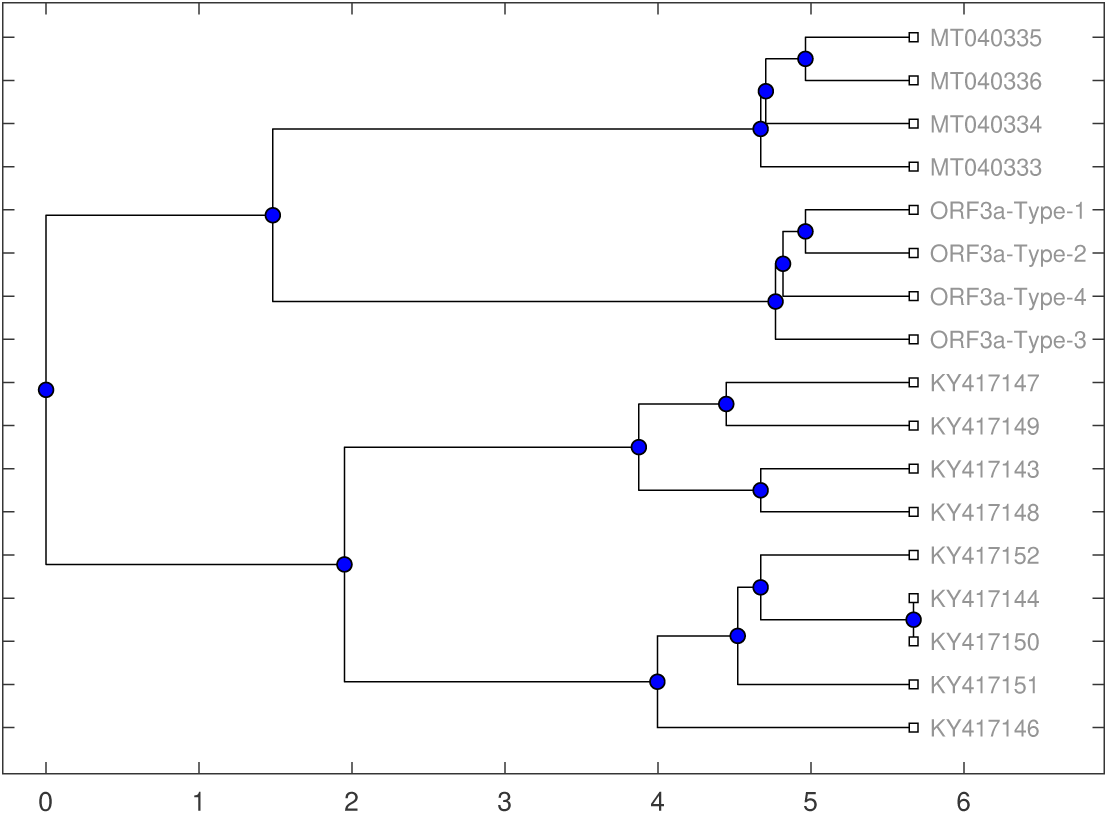
Phylogenetic relationships among the seventeen CoV genomes based on the frequency of amino acids in ORF3a proteins.

## 3. Conclusions

Among all the accessory proteins of SARS-CoV2, ORF3a is found to be very much important in playing virus pathogenesis as it possesses various mutations which are linked with that of the spike proteins. As mentioned, there are different mutations happened at various locations of the ORF3a gene of the SARS-CoV2 genomes of Indian patients and those mutations lead to alternation of amino acids. Among the mutations, the ORF3a-Type-3 and ORF3a-Type-4 mutations are restricted to only the Indian patients based in Ahmedabad so far it is identified. These mutations (Q to H, D to Y, S to L) are located near TRAF, ion channel, and caveolin binding domains respectively, suggesting that Type-3 and Type-4 might have effect on NLRP3 inflammasome activation. This unique non-synonymous mutations might affect the virulence of the virus and this needs a special attention from pathogenesis perspective by the medical scientists. A set of ORF3a genes of the Pangolin and Bat-CoVs were taken into consideration to investigate the evolutionary relationship from the phylogenies based on the nucleotides, dimers, codons and amino acids over the gene ORF3a across various genomes of CoVs. Based on conservations of nucleotide bases over the ORF3a genes, it is turned out that the ORF3a genes of four types of SARS-CoV2 and CoV-Pangolin are evolved from the ORF3a gene of the Pangolin CoV genome MT040333. It is worth noting that the ORF3a genes of Pangolin and Bat-CoV genomes are much more closer than that of SARS-CoV2, from the phylogenetic analysis of codon and amino acids conservations. From the molecular conservation analysis, it is emerged that the ORF3a genes across the seventeen genomes of SARS-CoV2 along with that of Pangolin and Bat-CoVs are co-evolved from the same origin.

## Author Contributions

SH conceived the problem. SH, PPC, PB and SSJ analysed the data and result. SH wrote the initial draft which was checked and edited by all other authors to generate the final version.

